# Well-tempered metadynamics calculations of free energy surfaces of benzothiadiazine derivatives in aqueous solution

**DOI:** 10.1101/2023.11.15.567238

**Authors:** Zheyao Hu, Jordi Marti

## Abstract

KRAS oncogenes are the largest family of mutated RAS isoforms, participating in about 30% of all cancers. Due to their paramount medical importance, enormous effort is being devoted to the development of inhibitors using clinical tests, wet-lab experiments and drug design, being this a preliminary step in the process of creating new drugs, prior to synthesis and clinical testing. One central aspect in the development of new drugs is the characterization of all species that can be used for treatment. In this aim we propose a computational framework based on combined all-atom molecular dynamics and metadynamics simulations in order to accurately access the most stable conformational variants for several derivatives of a recently proposed small-molecule, called DBD15-21-22. Free energy calculations are essential to unveil mechanisms at the atomic scale like binding affinities or dynamics of stable states. Considering specific atom-atom distances and torsional angles as reliable reaction coordinates we have obtained free-energy landscapes by well-tempered metadynamics simulations, revealing local and global minima of the free-energy hypersurface. We have observed that a variety of stable states together with transitions states are clearly detected depending on the particular species, leading to predictions on the behaviour of such compounds in ionic aqueous solution.

## 1. Introduction

KRAS proteins play a central role in a wide range of cellular processes such as signalling, survival, apoptosis or membrane trafficking, to mention a few[1, 2, 3]. They are binary switches between active GTP-bound and inactive GDP-bound states[4]. When active, they can interact with a big variety of effectors that control fundamental biochemical and biological processes such as EGFR[5, 6, 7] and they can also interact with downstream regulators such as MAPK[8, 9] or PI3K[10]. However, perhaps the most relevant role played by KRAS proteins is their crucial activity in about 30% of all known tumours[11, 12]. Due to this fact, many studies have been recently focused on the most important KRAS mutations (G12, V12, G13, Q61, etc.)[13, 14, 15, 16, 17] in order to find routes to the design and discovery of new effective clinical treatments and drugs. After near four decades of efforts, a major breakthrough was made ten years ago with the discovery of potential inhibitors of KRAS-G12C[18]. Later on, other new covalent inhibitors targeting KRAS-G12C such as AMG510[19, 20] and MRTX849[21] were proposed. Finally, back in 2021 a drug called “sotorasib” was finally approved by the U.S. Food and Drug Administration as the first clinical treatment of KRAS-G12C based cancers[22, 23]. In the case of KRAS-G12D, a variety of different several strategies has already been proposed: use of indole-based inhibitors targeting the well-known Switch-II pocket[24, 25], targeting the piperazine-based compound TH-Z835 for ASP12 residue[26], targeting the site PRO110[27] or a specific three cyclic peptide[28] among others. Recently, mutations G12R, G12S and G12V of KRAS were also studied and some new inhibitors were proposed[29, 30, 31, 32, 33, 34]. Nevertheless, despite the progress targeting oncogenic KRAS-G12D still remains elusive and, up to the best of our knowledge, no other therapeutic agents have been clinically approved.

Detailed information on atomic interactions and local structures at the all-atom level of KRAS are of crucial importance in the design of potential compounds able to target oncogenic KRAS[35], given the difficulty to access nanoscale length and time in experimental setups. In a previous work by means of microsecond time-scale molecular dynamics simulations, it was reported that the mutation of GLY12 site in KRAS-G12D is able to trigger a particular change of the interaction patterns between Mg^2+^ and KRAS. Furthermore, two druggable exclusive dynamic pockets on the KRAS-G12D-GDP-Mg^2+^ ternary complex surface were revealed[36]. Getting advantage of the full knowledge of the structural characteristics of these two dynamic pockets, a specific inhibitor (DBD15-21-22, a derivative of benzothiadiazine (DBD)[37, 38], was designed *in silico*. Such compound can specifically target the KRAS-G12D oncogene, stabilizing the inactive state GDP-KRAS-G12D complex. One particular property that remains to be analyzed is the multidimensional free-energy (hyper)surface (FES) of DBD15-21-22 in solution and when associated to KRAS-G12D. In order to advance in this direction, we report in the present work a series of free-energies obtained in aqueous solution for the original DBD15-21-22 compound and four of its derivatives. To do so, we have employed all-atom molecular dynamics (MD) and the FES has been obtained by well-tempered metadynamics (WTM) simulations[39], in order to acquire precise information on the preferential conformations. These two classical simulation tools have revealed valuable information on a wide variety of systems such as small-drugs at membrane interfaces[40] or in the mechanism of anchoring of KRas-4B[41, 39]. For the present work, we will use a combined protocol so that we can select from MD the most convenient collective variables (CV), which will be the cornerstone of the present and future WTM simulations.

## 2. Methods

### 2.1. System Preparation and MD Simulations

MD simulations are a highly successful tool for the modeling and simulation of atoms, molecules and small solutes in bulk and at interfaces[42, 43, 44, 45, 46], including its recent strong impact on drug discovery[47]. MD is also able to render microscopic properties connected directly to experiments, such as all sorts of neutron diffraction and spectroscopic data (X-ray, infrared and Raman spectroscopy, etc.)[48]. For the accurate modeling and simulation of biosystems[49, 50, 51, 52], several reliable force fields such as AMBER, OPLS and CHARMM[53, 54, 55, 56, 57] have been developed. These works include also investigations on the behaviour of small molecules in bulk an in solvated phospholipid membranes[58, 59, 60, 61, 62, 63].

From a general perspective, in a system with multidimensional reaction coordinates usually several stable states (bound configurations) are separated by high free energy barriers corresponding to transition states of the system[64, 65], making it difficult for MD simulations to sample them adequately[66]. Nevertheless, free energy calculations using enhanced sampling techniques provides a method to address the problem, acquiring meaningful understanding of the passive diffusion phenomena of small solutes over a barrier, which requires a detailed view of the underlying free energy surface. Once the FES is described, minimum free energy paths can be traced[67] and transition states can be located[68]. To the best of our knowledge, the free energy landscapes of small-solutes acting as potential drugs in solution is still mostly unknown. This is partially due to the difficulty of applying appropriate sampling techniques to address the problem, and also because the determination of the proper collective variables is always a difficult challenge[69, 70] (see Section 2.2).

We conducted a series of MD simulations of single prototype drugs derived from a benzothiadiazine species, as reported previously[71, 36]. The derivatives from DBD15-21-22 were embedded inside an aqueous ionic solution of sodium chloride and magnesium chloride (0.15 M concentration) at 310.15 K and the pressure of 1 atm in all cases. All MD inputs were generated using the CHARMM-GUI solution builder[72, 73, 74] and the CHARMM36m force field[75] was adopted for all interactions. In particular, the water model employed was TIP3P[76] in order to achieve full equilibration and collection of meaningful properties we extend the total MD runs up to the time length of 4.8 *μ*s. The full system is shown in Figure1.

**Figure 1.**
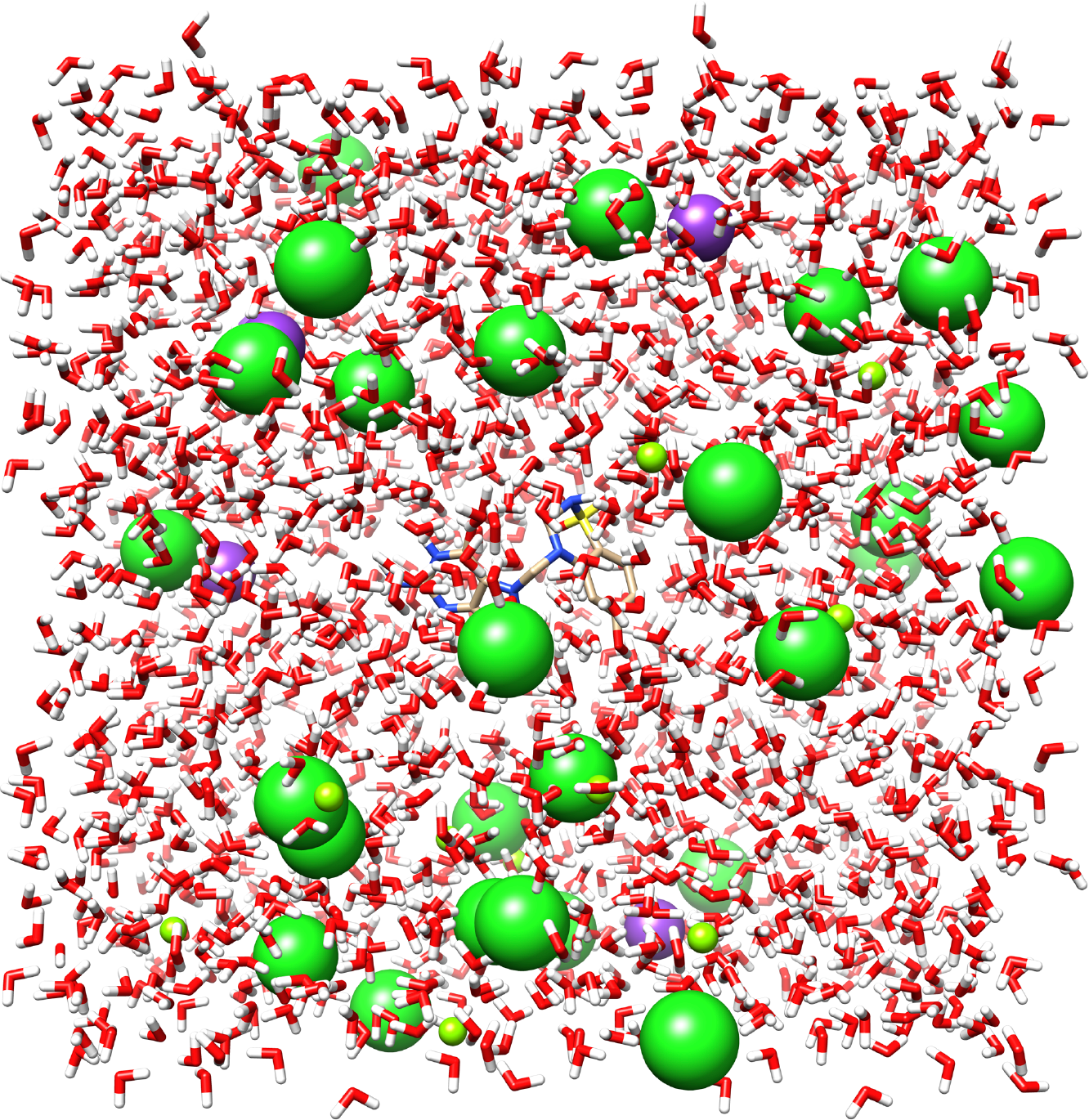
DBD15-21-22s in water box along with NaCl and MgCl_2_ at 310.15K and 1 atm.

### 2.2. Well-tempered Metadynamics

A wide variety of methods has been proposed to handle the problem of computing free energy landscapes in multidimensional quantum or classical systems, such as hybrid quantum mechanics/molecular mechanics methods[77], transition path sampling[78, 79, 80, 81, 82, 83], or others based on multidimensional order parameters like adaptive biasing force[84], multi-state empirical valence bonds[85, 86] or umbrella sampling methods[87, 88]. Others, such as density functional theory molecular dynamics[89] or calculations of potentials of mean force[90] based on reversible work methods[60] use simple, one-dimensional order parameters such as the radial distance between atomic sites. However, when other degrees of freedom orthogonal to the biased one are of importance, the obtained FES may contain errors that could easily lead to wrong conclusions. In the present work we have employed WTM, a method able to efficiently explore free energy surfaces of complex systems using multiple reaction coordinates, very successful for a wide variety of complex systems[91, 66, 92, 93, 94, 95]. The main advantage of WTM is its suitability for a wide variety of systems, including model cell membrane systems with attached small-molecules and proteins[68, 39, 63], recently developed in our group.

After equilibrating systems composed by one DBD15-21-22 derivative in aqueous solution from the previous MD run, we switched to run another 0.8 *μ*s of WTM simulations to perform Gibbs free-energy calculations, starting from the last configuration of MD simulations. Long well-tempered metadynamics simulations were performed using the joint GROMACS/2021-plumed-2.9.0 tool[96, 97]. The isothermal-isobaric NPT ensemble at temperature 310.15 K and pressure 1 atm was adopted in all cases. Periodic boundary conditions in the three directions of space were also considered. The CV adopted in the WTM simulations will be discussed in full details below (see Section 3).We should point out that determining and using more than two CVs in a WTM simulation is a hard challenge from the computational point of view and would produce a four-dimensional FES, unsuitable to deal with. A very usual choice is to consider one or two CVs[98]. From our experience, the use of two CVs produces a complete enough description of the FES and it is the optimal way to proceed. Later, it is a usual procedure to project the free energies onto one single coordinate, integrating out the contribution of the second CV.

The values for the parameters of the metadynamics simulations[99] are listed in Table 1.

**Table 1.**
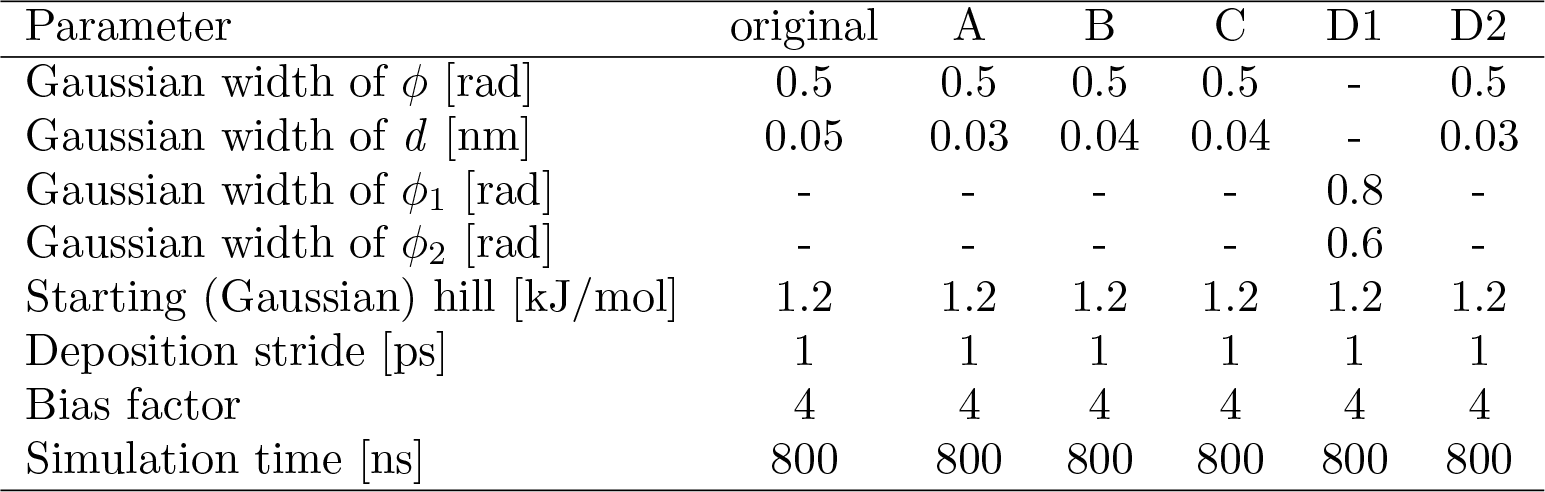
WTM simulation parameters.

**Table 2.**
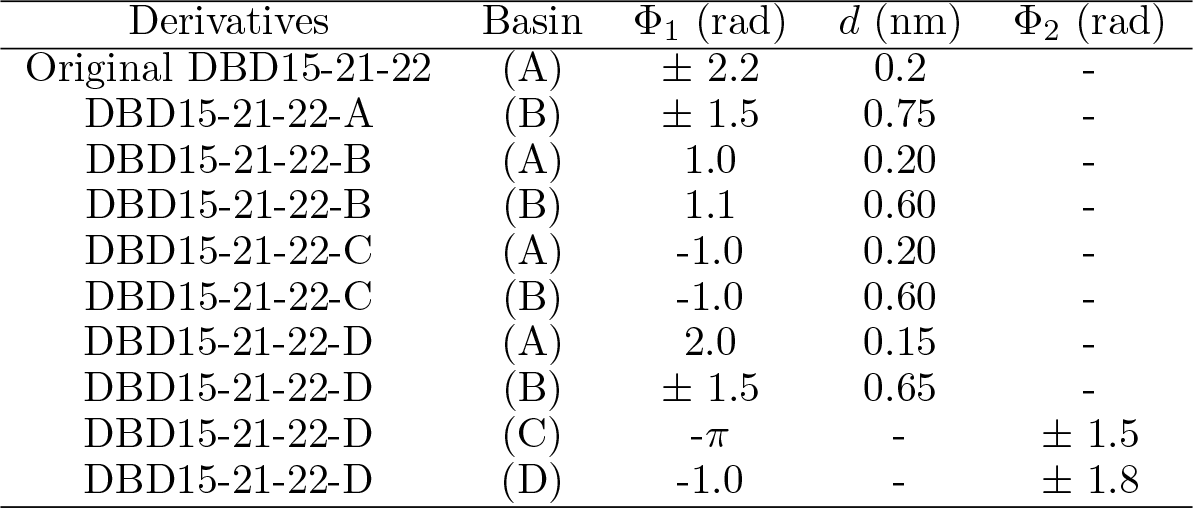
Location of the stable basins of all derivatives. All distances are given in nm and angles are given in rad.

| Derivatives | Basin | $\Phi_1$ (rad) | $d$ (nm) | $\Phi_2$ (rad) |
| --- | --- | --- | --- | --- |
| Original DBD15-21-22 | (A) | $\pm 2.2$ | 0.2 | - |
| DBD15-21-22-A | (B) | $\pm 1.5$ | 0.75 | - |
| DBD15-21-22-B | (A) | 1.0 | 0.20 | - |
| DBD15-21-22-B | (B) | 1.1 | 0.60 | - |
| DBD15-21-22-C | (A) | -1.0 | 0.20 | - |
| DBD15-21-22-C | (B) | -1.0 | 0.60 | - |
| DBD15-21-22-D | (A) | 2.0 | 0.15 | - |
| DBD15-21-22-D | (B) | $\pm 1.5$ | 0.65 | - |
| DBD15-21-22-D | (C) | $-\pi$ | - | $\pm 1.5$ |
| DBD15-21-22-D | (D) | -1.0 | - | $\pm 1.8$ |

**Table 3.**
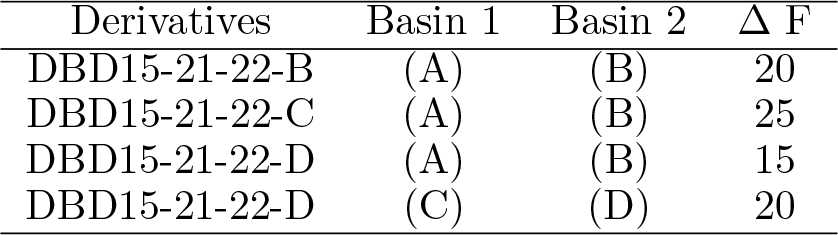
Estimation of the free energy barriers Δ*F* between stable states. All values are given in kJ/mol.

| Derivatives | Basin 1 | Basin 2 | $\Delta F$ |
| --- | --- | --- | --- |
| DBD15-21-22-B | (A) | (B) | 20 |
| DBD15-21-22-C | (A) | (B) | 25 |
| DBD15-21-22-D | (A) | (B) | 15 |
| DBD15-21-22-D | (C) | (D) | 20 |

## 3. Results

In a previous work, a prototype of a drug candidate for KRAS-G12D was proposed[36]. This species, called DBD15-21-22 (see Figure 4 of this reference), is a derivative of DBD[37, 38] able to target a given pocket in the inside of KRAS-G12D oncogene (see Figure 3 of this reference), fixing GDP inside such pocket and favoring the inactivity of KRAS by difficulting its binding to GTP and its capacity of signalling when anchored to the cell membrane inside the cytoplasm. As indicated above, the FES of DBD15-21-22 has not been reported yet, so that the free-energy barriers to be surmounted for the small-molecule in order to access the drugging pocket are not known. This information would be of crucial importance in order to evaluate the reliability of the drug when ingested as a pharmaceutical candidate. In order to access the FES, it is necessary to know the best reaction coordinates (or CV) to be employed in its calculation. In the case of metadynamics, the choice of such coordinates is of paramount importance to obtain a clear and well defined FES. In the present work, we will evaluate the best CV candidates for DBD15-21-22 in aqueous ionic solution, in order to use such variables in the forthcoming computation of the FES of DBD15-21-22 when locked inside the KRAS-G12D surface.

Firstly, we will describe the prototype drug DBD15-21-22 and its four derivatives and secondly we will propose two different sets of CV, evaluating their reliability by computing the FES in each case and corroborating the convergence of the WTM simulations. Finally, the physical meaning of the results will be discussed, indicating the most suitable choice for the CV in each case.

### 3.1. Derivatives of GDP bound KRAS-G12D able to block its active state

A detailed study of a DBD derivative able to block the GDP guanosine inside its natural groove at KRAS-G12D was recently reported[36]. However, in order to improve the feasibility for the synthesis of the proposed DBD15-21-22 compound, four structural variations of the original small-molecule were proposed by Kessler and co-workers[100]. The chemical structures of the variations are reported in Figure2. They include the original prototype drug (DBD15-21-22) as well as the replacement of a nitrogen by a carbon atom (DBD15-21-22-A), the change in a chemical bond (double to single, DBD15-21-22-B), isomeric version of DBD15-21-22-B (DBD15-21-22-C) and the extension of a particular hexagonal structure (DBD15-21-22-D).

**Figure 2.**
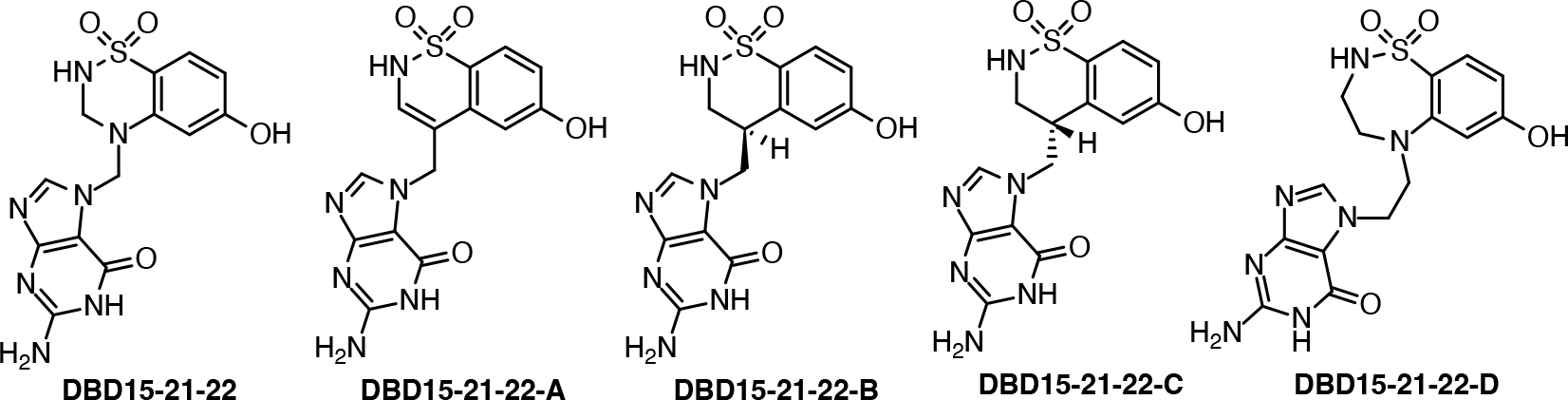
Chemical structures of DBD15-21-22 derivatives considered in this work.

### 3.2. Well-tempered metadynamics simulations: Collective variables

The enhanced sampling method WTM applies a time-dependent biasing potential along a set of CV by adding a Gaussian additional bias potential to the total potential in order to overcome barriers larger than *k*_*B*_*T*, with *k*_*B*_ being Boltzmann’s constant and *T* the temperature. In the method, sampling is performed on a selected number of degrees of freedom (*s*_*i*_), i.e. the set of CV. For each degree of freedom, the biased potential *V* (*s*_*i*_, *t*) is a dynamical function constructed as the sum of Gaussian functions[101, 99, 102]:

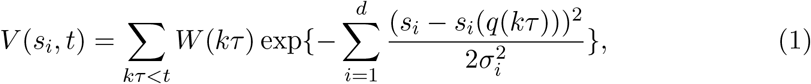

where *k* is an integer, *τ* is the Gaussian deposition stride, *W* (*kτ*) is the height of the Gaussian and *σ*_*i*_ is the width of the Gaussian for the i-th CV. The choice of CV is of central importance. Good CV must be able to distinguish between initial and final states, while properly describing all intermediate states. Usually CV are taken as relative distances between atomic sites of the small-molecule or some particularly relevant internal angles in the molecule. In the present work, we have taken a specific approach similar to previous works[68, 103] where distances combined with torsional angles or a pair of dihedral angles were chosen.

After a thorough selection from distances and angular coordinates, the two CVs selected to perform the 2D WTM calculations are (see Figure3): (1) distances *d* between the marked atoms (in red) and (2) the dihedral angles (*φ*_*i*_) formed by the groups of four atoms marked in blue in all cases. We additionally considered the combination of two torsional angles in the case DBD15-21-22-D, marked in blue and green. Single angular variables were not producing suitable, i.e. well defined FES. Nevertheless, the selection of the CV can only be assessed and evaluated when the FEs is computed. However, it is possible to have an initial indication of the quality of each CV pair by observing how the phase space of the two coordinates is sampled along a full WTM trajectory. The results for each CV are reported in the Section 3.3.

**Figure 3.**
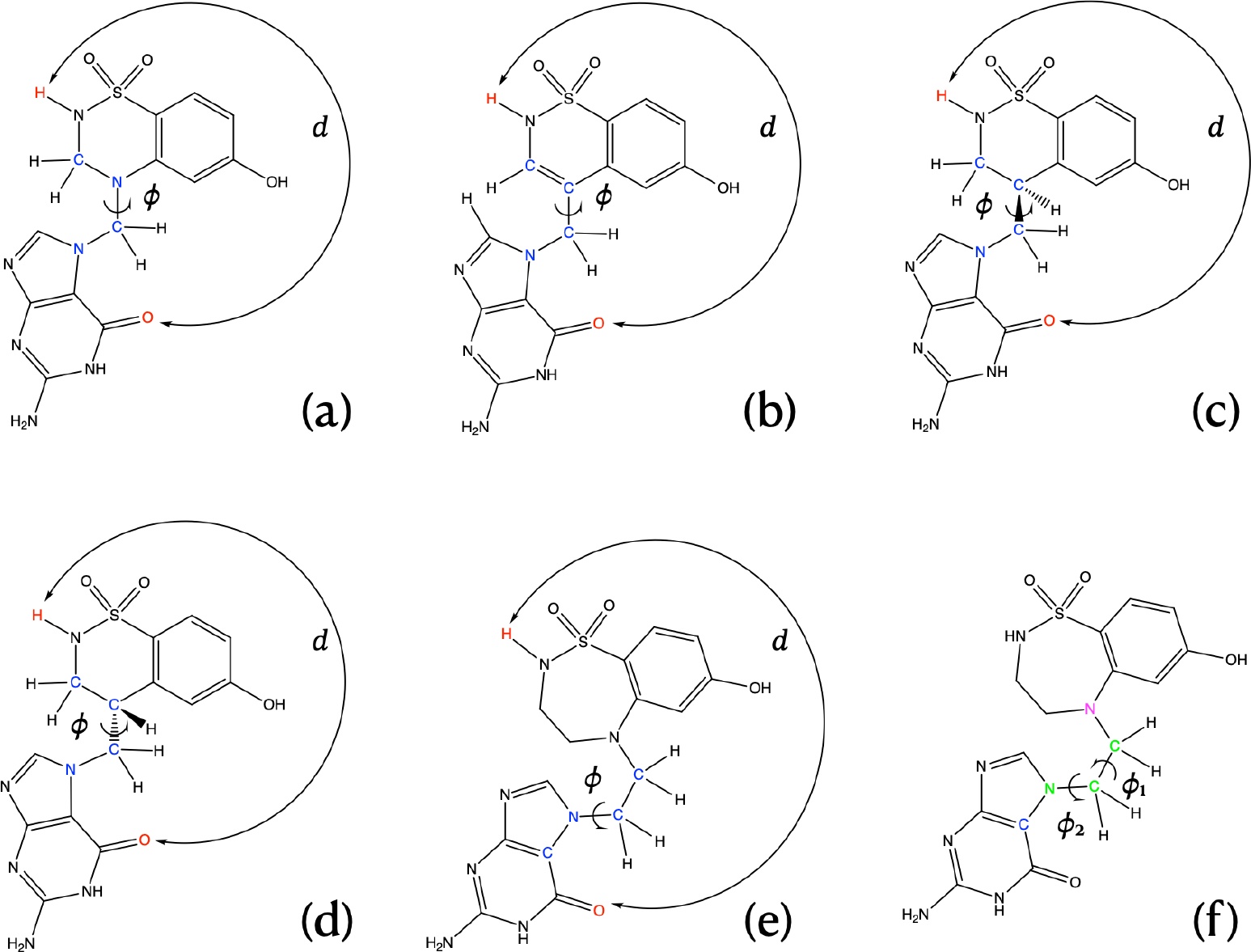
CV sets employed in the present work: (a) Used for DBD15-21-22; (b) for DBD15-21-22-A; (c) for DBD15-21-22-B; (d) for DBD15-21-22-C; (e) for DBD15-21-22-D; (f) for DBD15-21-22-D

### 3.3. Well-tempered metadynamics simulations: two-dimensional free-energy landscapes

Two dimensional (2D) free-energy landscapes are represented in Figure 4 for the cases of the original compound DBD15-21-22 and derivatives DBD15-21-22-A and DBD15-21-22-B, whereas the compounds DBD15-21-22-C and DBD15-21-22-D are reported in Figure 5.

**Figure 4.**
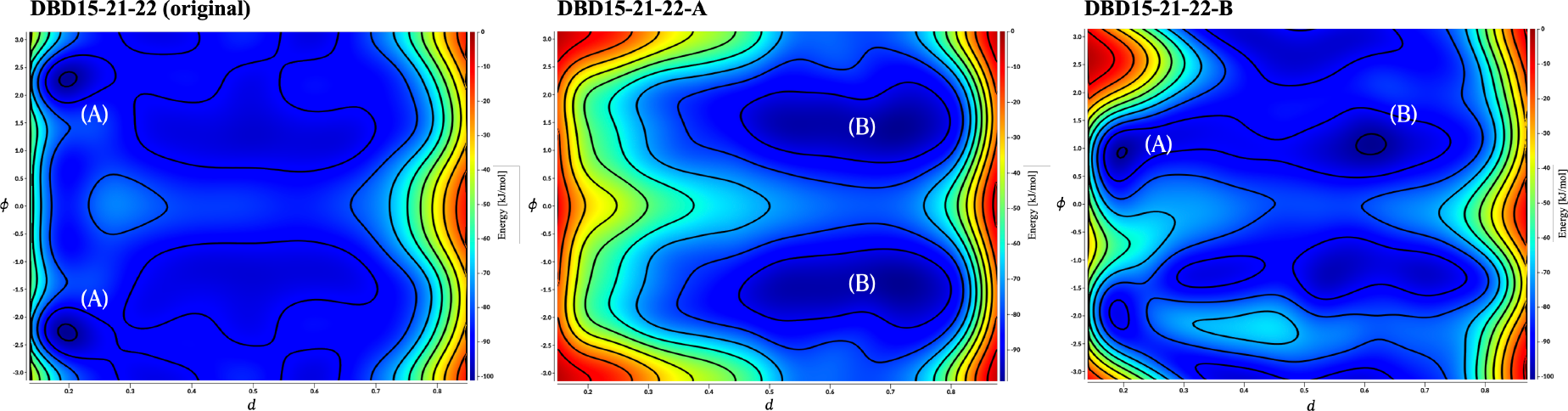
2D free-energy landscape F(*CV*_1_, *CV*_2_) (kJ/mol) for the compounds; (a) DBD15-21-22; (b) DBD15-21-22-A and (c) DBD15-21-22-B. All distances are given in nm and angles are given in rad.

**Figure 5.**
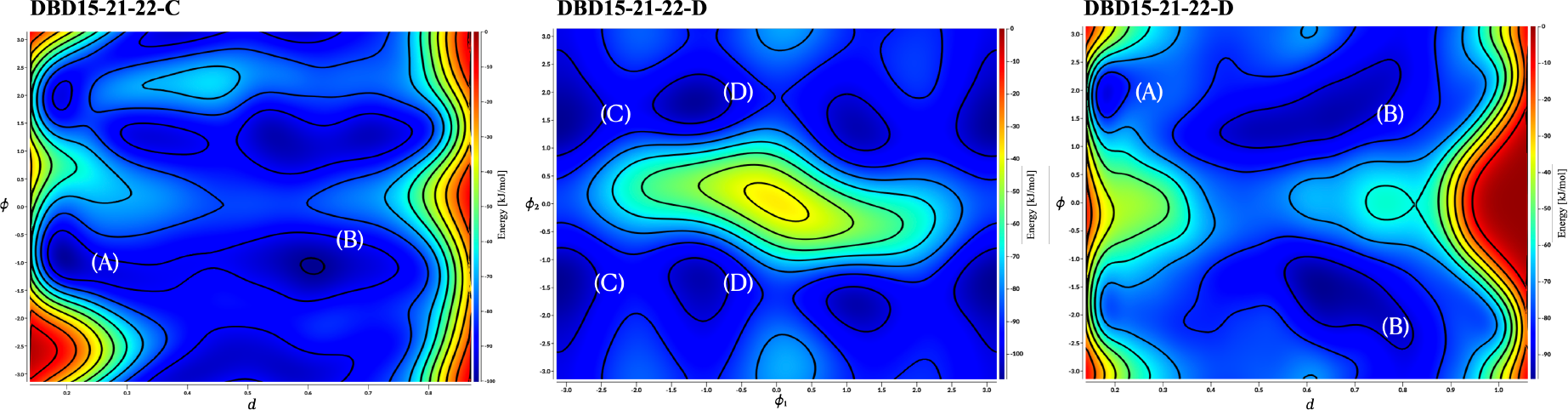
2D free-energy landscape F(*CV*_1_, *CV*_2_) (kJ/mol) for the compounds; (d) DBD15-21-22-C; (e) DBD15-21-22-D with dihedral angles as CV and (f) DBD15-21-22-D, with specific distance and torsional angle as CV. All distances are given in nm and angles are given in rad.

We can observe that two main basins (A, B) are well defined in all cases. In addition, for the derivative DBD15-21-22-D two different basins arise (C,D) when we use two dihedral angles as CV. We have only indicated the basins which are clearly seen from the FES, assuming that even with well converged runs the definition of the FES requires a huge amount of statistics, probably of the order of 5 *μ*s, far from the actual possibilities of our lab. The coordinates of the stable state basins are reported in Table2.

As a general concept, the stable basins in the FES indicate the most probable configurations of the system in equilibrium. As a matter of fact, WTM is able to reveal those stable configurations in such a way that the free energy barriers that the system needs to overcome to shift between stable states can be estimated with a high degree of accuracy. In this work, we can distinguish at least one basin for each derivative proposed. In the cases of the original DBD15-21-22 species and its first derivative DBD15-21-22-A only one stable state can be clearly observed given the CV set considered, whereas in the cases of derivatives DBD15-21-22-B, DBD15-21-22-C and DBD15-21-22-D, at least two basins are well defined.

In the particular case of DBD15-21-22-D, we have employed two sets of CV, the first one with the standard choice of one distance and one torsional angle and the second one using two dihedrals. The minima (A)-(B) at the FES correspond to two different distances *d* (0.15, 0.65 nm) and one torsional angle between 1.5-2 rad. Besides, the minima obtained from the combination of the two dihedrals (C) and (D) revealed a torsional angle of about 1.5-1.8 rad as expected and, in addition, two corresponding stable dihedrals around -1 and -*π* rad which were not detected in the previous calculation using distance-single dihedral CV. This suggests that: (1) the choice of the CV is truly relevant and can help to uncover stable configurations difficult to find using standard variables: (2) the derivative DBD15-21-22-D possesses a richer free energy landscape with at least three different stable basins.

The energy barriers between stable basins will indicate the amount of free-energy required for the DBD derivatives to exchange their conformations between two stable configurations, as it was described in Ref. [103]. We are reporting in Table3 estimations of the barriers associated to transitions between stable states for each derivative, although a more precise is left to Section 3.4.

Being this a rough estimation based in the two dimensional FES reported in Figures 4 and 5, we can only extract qualitative information: all five species analysed in the present work require moderate energies between 15-25 kJ/mol to exchange their stable configurations, indicating that all compounds will likely be able to bind to a drugging pocket in a similar manner. However, only specific WTM simulations of the derivatives inside such pockets would reveal the precise energy barriers to surpass for a stable binding. The values reported here are of the same order of magnitude as those found by Jämbeck et al.[40] or by Lu et al.[68] in the case of small-molecules close to the surface of model cell membranes.

### 3.4. One-dimensional free-energy profiles

From the 2D free energy landscapes obtained from WTM simulations (Figs. 4 and 5) it is possible to obtain a one-dimensional (1D) free-energy profile where one single CV is considered and the second one has been integrated out. This calculation along one single CV (*s*_*i*_, *i* = 1, 2), allows us to directly compare free-energy barriers to experimental findings, as it was pointed out by Jämbeck et al.[40]. These authors pointed out that a direct route to connect FES to experiments is normally cumbersome, since information about binding modes in solutes obtained from experiments is normally not available. Then, it is possible to use standard binding free energies Δ*G*_*bind*_ determined from experiments as an indirect measurement of the accuracy of computed free energy barriers. In the present work, 1D free-energy profiles F(*s*_1_) (equivalent to the Δ*G*_*bind*_ mentioned above) can be obtained as [40, 104]:

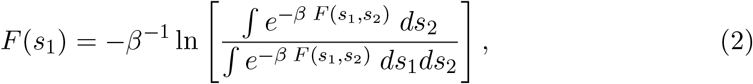

where *s*_1_ and *s*_2_ are the CV, *β* = 1*/*(*k*_B_*T*), *k*_B_ is the Boltzmann constant and T is the absolute temperature. This means that all possible paths for the CV labelled as *s*_2_ have been integrated out and averaged. In the present case, the results of Figures 6 and 7 reveal a series of free energy barriers around the stable basins.

**Figure 6.**
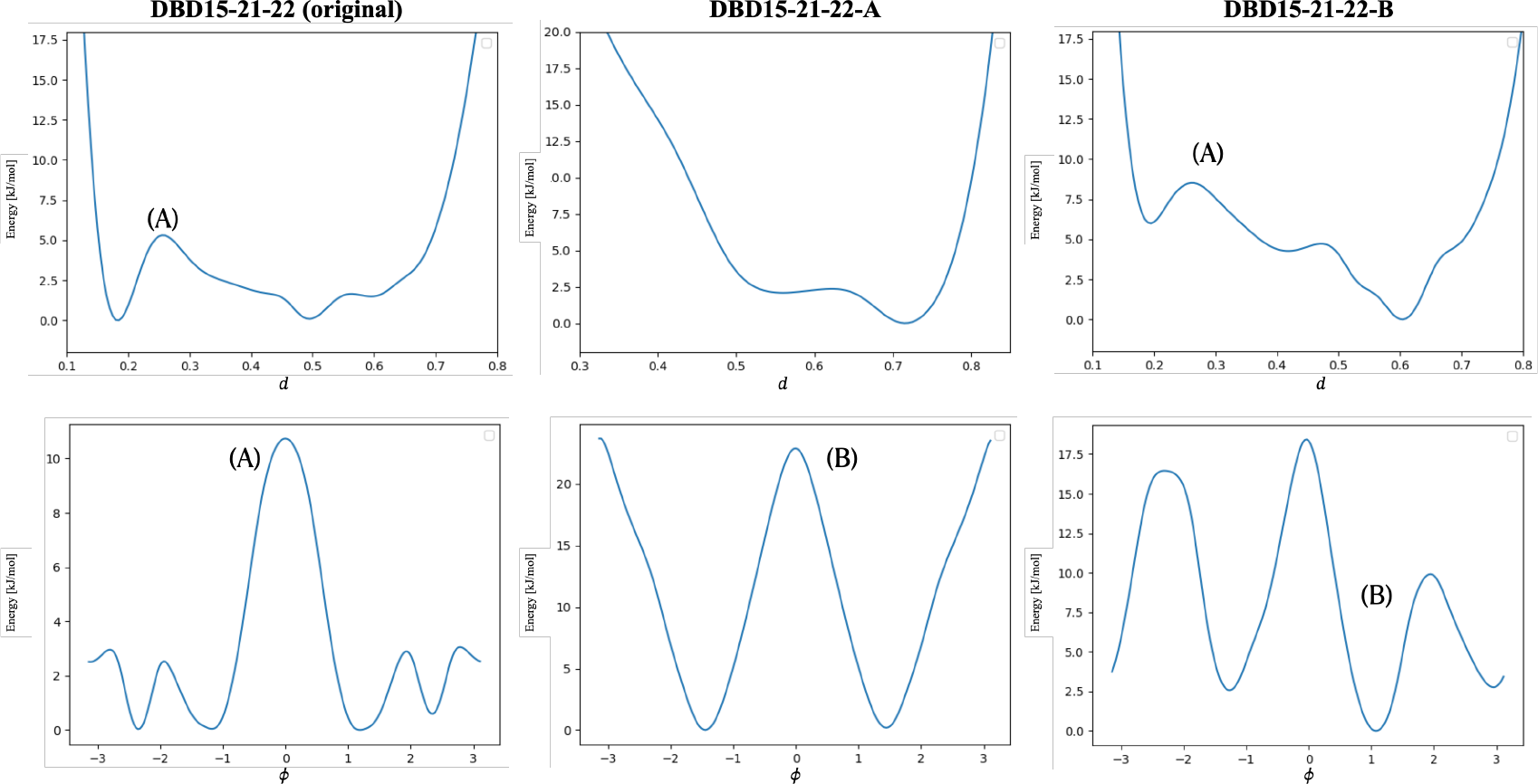
1D integrated free-energy profiles for two CV. Top row correspond to the distance CV *d*, whereas bottom row correspond to the CV dihedral angles defined in Figure 3 for each derivative. Basin regions marked with same labels (A, B) as in Fig.4. In order to directly compute the height of free-energy barriers, absolute minima are set equal to zero. Free-energy profiles generated by the PLUMED2 tool.

**Figure 7.**
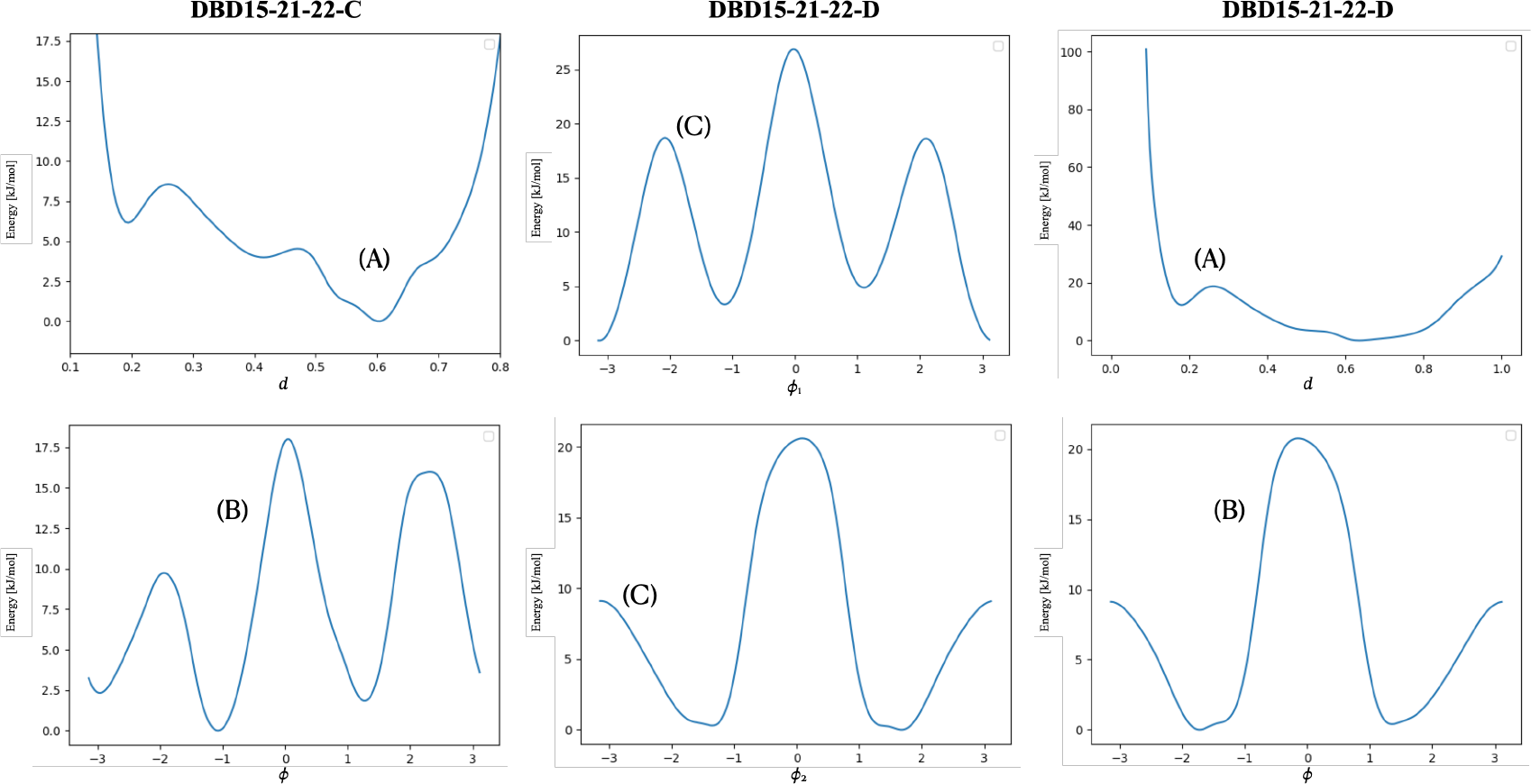
1D integrated free-energy profiles for two CV. Basin regions marked with same labels (A, B, C, D) as in Fig. 5. In order to directly compute the height of free-energy barriers, absolute minima are set equal to zero. Free-energy profiles generated by the PLUMED2 tool.

We have obtained quasi-symmetrical profiles for the dihedral angle’s coordinates and clear asymmetry for all free energies dependent of *d*, indicating the existence of overall two equivalent orientations for each coordinate, but distinguishable distributions for the distances. As a general fact, barriers for the free energies dependent of distances (computed after integration of the torsional angles) are much lower than those obtained from free energies dependent of dihedrals. The former are in the range of 5 kJ/mol the largest, and the latter correspond to values between 15 and 25 kJ/mol. These numbers clearly indicate that free energy changes should be attributed to fluctuations of the torsional angles rather than to changes in coordinate *d*.

Up to the best of our knowledge no solvation free energies of benziothiadiazines have been previously reported. Interestingly, Zhou et al.[105] reported results of meta-dynamics simulations of KRAS proteins in plasma membranes using angular collective variables, finding barriers of the order of 12 kJ/mol, quite close to the ones reported in the present work. In a different fashion, Chen et al.[106] computed binding free energies of hypoxanthine bound to guanine riboswitches in water solution, finding values in the range of 2-8 kcal/mol, again similar to those reported in the present work.

### 3.5. Convergence of the WTM simulations and free-energy profiles

Now that 1D profiles have been defined and analysed, we can complete our study of the convergence of the WTM simulations reported here. Concerning the reliability of the simulations reported in the present work, we can use several criteria. Three of the most efficient and reliable are: (1) the time evolution of the fluctuations of the hills of the biased potential along the full trajectory; (2) the time cumulative average of 1D free energy profiles as defined above (Section 3.4) and (3) considering the so-called block analysis of the average error along free-energy profiles as a function of the block length.

The evolution of the hills is presented in Figure 8. We observe a overall decreasing behavior, eventually including low-height spikes. In all cases, the height of the biased potential decreased accordingly along the simulation runs. In all cases, a quasi-flat profile is already seen around 500-600 ns. This is a clear indication of the convergence of the WTM runs.

**Figure 8.**
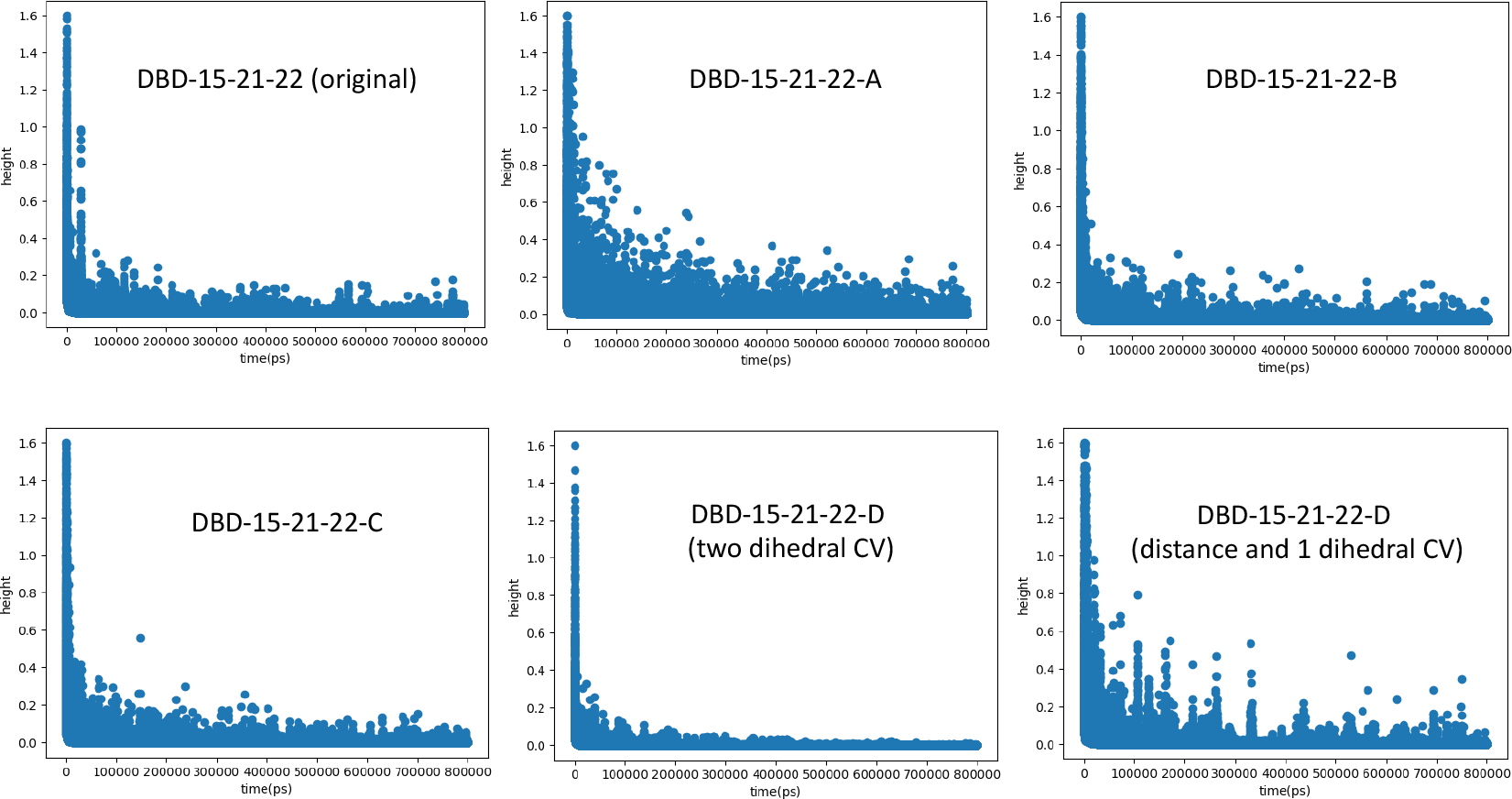
Height of the hills of the Gaussian-biased potential as a function of time. All profiles correspond to the WTM simulations of each derivative. In the case of DBD15-21-22 the two sub-figures correspond to the two different WTM simulations using different sets of CV, as indicated in Figure 3.

From the results of Figure 9 we can see that after long cumulative time lengths, the differences between a profile and the one immediately before are very small (up to 2.5 kJ/mol) and lead us to fully converged free energies for the two CV considered in the present study. In order to limit the number of cases presented, we have considered the cumulative profile of *d* for all derivatives except DBD15-21-22-D and the two angular CV in the case of DBD15-21-22-D. However, in all cases very well converged profiles can be observed around 500 ns, regardless of the type of CV.

**Figure 9.**
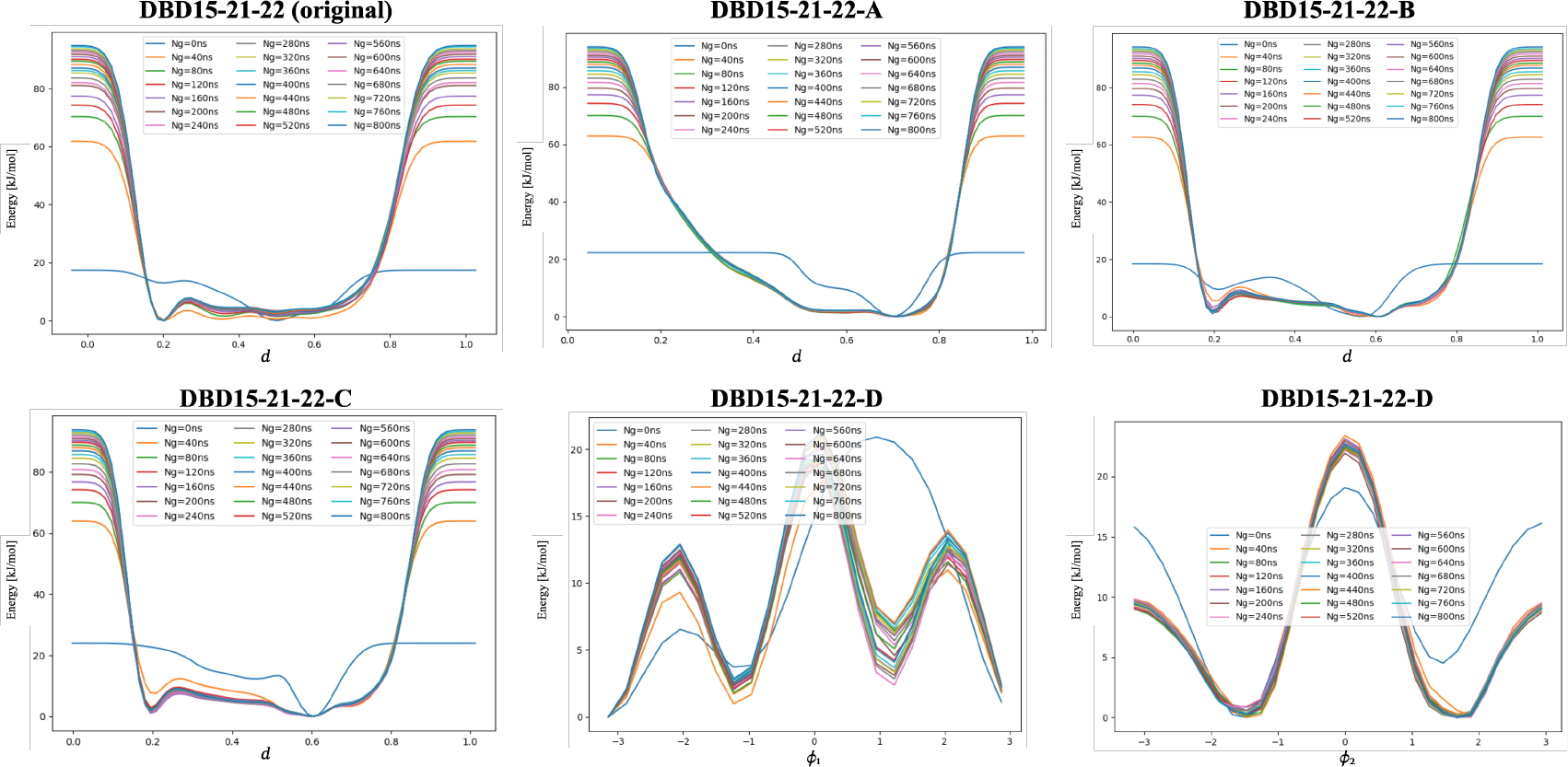
Selected time cumulative free energy pro_les for each derivative in the full range of the WTM simulations.

Finally, from the results of Fig. 10, where we report the size of the average error in the free-energy profiles calculated from two different sets of well-tempered meta-dynamics simulations as a function of the block size. The initial 25000 steps of all metadynamics trajectories were discarded. As expected, the errors increase with the block length until they reach a plateau in all cases. The average errors tend to stabilise around 0.1 kJ/mol for all cases considered. For this calculation, scripts provided by the PLUMED project[96, 97] have been employed.

**Figure 10.**
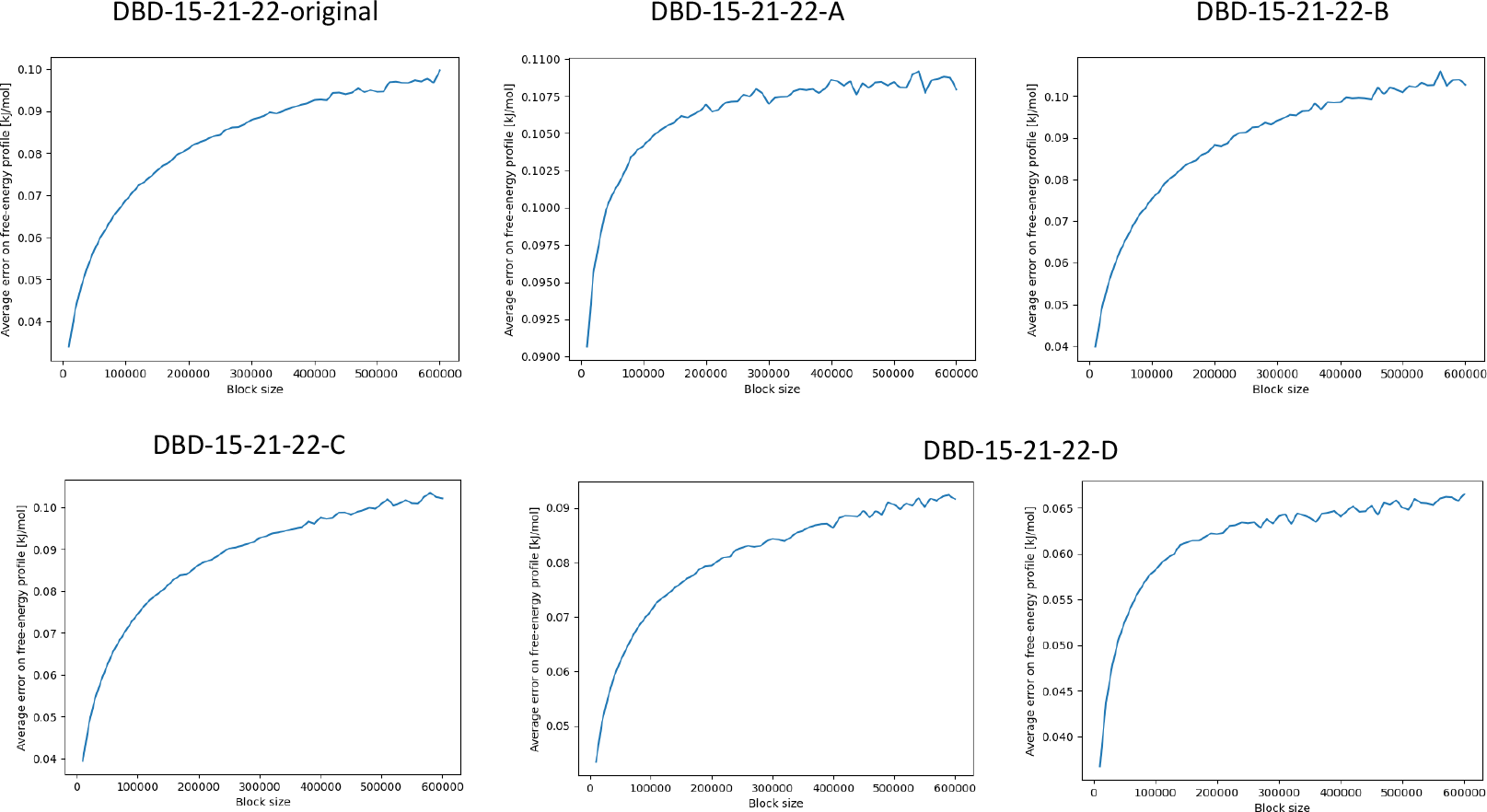
Block-size averages. Original DBD15-21-22: dihedral angle; DBD15-21-22-A: distance; DBD15-21-22-B: distance; DBD15-21-22-C and DBD15-21-22-D: dihedral angle.

## 4. Conclusions

KRAS-G12D proteins are small GTPases with the ability to regulate cell growth, differentiation and survival. They are related to cancers such as in lung, colon and pancreas. Even though great efforts have been focused on the structural and pharmaceutical aspects of KRAS-G12D, efficient inhibitors are still missing. In the present work, we have considered a series of prototype drugs derived from a compound (DBD15-21-22) reported recently[36] and we have evaluated the free energy landscapes of all the derivatives in aqueous solution in NaCl and MgCl_2_ at the human body concentration, at 310.15 K and at the fixed pressure of 1 atm. The present study is previous to the analog where the five species considered here will be located inside the oncogene KRAS-G12D and their free energy profiles will be computed. As a general fact, free energy landscapes of inhibitors in solution are very much unknown since, up to date, only a few small-molecule analogs have been simulated in the vicinity of a lipidic membrane. In this work, we have conducted MD and WTM simulations of five derivatives of a prototype compound based on benzothiadiazine and obtained the corresponding FES, considering in all cases a specific pair of reaction coordinates or collective variables. Our simulations reached the scale of 0.8 *μ*s. In all cases the convergence of the WTM simulations has been clearly proved.

With the help of well-tempered metadynamics simulations we have calculated 2D FES and the 1D profiles after integrating out one CV. Our results indicate that each system has at least one stable state, although in the cases of DBD15-21-22-B and DBD15-21-22-C we have obtained two stable basins and in the case of DBD15-21-22-D we can identify up to three stable states. These states are the signature of the equilibrium configurations for each system. Furthermore, the stable state of original DBD15-21-22 found in this paper matches the state previously obtained through classical MD simulations, see Figure 5 of reference[36]. The WTM simulations have given us quantitative information on the energy barriers that must be surmounted by DBD15-21-22 and its derivatives in order to change its orientation and eventually its internal structure. The analysis of the 2D and 1D free energy profiles revealed barriers between stable states of the order of 15 to 25 kJ/mol, numbers which are in good qualitative agreement with other values reported in the literature, related to the solvation of small-molecules and drugs near model cell membranes or binding of species such as hypoxanthine-guanine in aqueous solution. These are quite large barriers and they suggest that it will require long periods of time and eventual increase of temperature or pressure for the derivatives to be able to access its second preferential stable orientation from a primary one.

## Acknowledgement(s)

We thank Dirk Kessler from Boheringer Ingelheim for the suggestion of the four derivatives of DBD15-21-22 employed in this work.

## Disclosure statement

No potential conflicts of interest

## Funding

We thank financial support provided by the Spanish Ministry of Science, Innovation and Universities. This publication is a part of the I+D+i project with reference PID2021-124297NB-C32, founded by MCIN/AEI/10.13039/501100011033 and “FEDER Una manera de hacer Europa-A way of making Europe” by the “European Union NextGenerationEU/PRTR”. Zheyao Hu is a Ph.D. fellow from the China Scholarship Council (grant 202006230070). J.M. thanks the *G*eneralitat de Catalunya for the support through the grant *G* rup de Recerca SGR-Cat2021 Condensed, Complex and Quantum Matter Group reference 2021SGR-01411 and to the Polytechnic University of Catalonia-Barcelona Tech through the funding AGRUPS. Computational resources provided by the Barcelona Supercomputing Center, project BCV-2023-2-0004 are also acknowledged.

## Notes

### Competing Interest Statement

The authors have declared no competing interest.

